# Agentic AI for Structural Elucidation and Discovery of Drug Metabolites from Mass Spectrometry Data

**DOI:** 10.64898/2026.06.23.734138

**Authors:** Xianghu Wang, Abubaker Patan, Haoqi Nina Zhao, Vincent Charron-Lamoureux, Yourae Shin, Daniel Petras, Yuhui Hong, Benjamin P. Bowen, Trent R. Northen, Pieter C. Dorrestein, Mingxun Wang

## Abstract

The majority of chemical signals detected in public metabolomics repositories remain structurally undefined. Large language models (LLMs) are probabilistic systems whose capacity to generate outputs beyond their training data, which can cause hallucinations, makes them also potentially suited to hypothesize structures for molecules that have never been described. We aimed to build a system that could harness this LLM generative capacity combined with domain specific tools/framework to constrain hallucination and produce validated discoveries. We developed a GNPS2 agentic AI system that interprets LC-MS/MS data by integrating spectral alignment, molecular formula inference, rule-based structural enumeration, machine learning-based spectrum prediction, and translates natural language hypotheses from domain experts into dynamically generated analytical workflows. We demonstrate the annotation of unknown drug metabolites from public data guided by chemical hypotheses. The agent predicted, and we experimentally confirmed, a phosphorylated hydroxyzine, an acetaminophen-p-coumaric acid ester, and identified two new oxidative ibuprofen-carnitine conjugates from public repositories. These results demonstrate that LLM-driven agentic reasoning, when combined with domain expertise, can indeed generate experimentally testable structural hypotheses for previously uncharacterized metabolites leveraging pan repository data.

## Introduction

Large language models (LLMs) integrated into agentic AI frameworks (e.g., Claude Code, Codex) are rapidly transforming how analytical workflows are designed and executed. Rather than relying on fixed computational pipelines, analyses can increasingly be constructed on demand to address specific scientific questions. Early LLMs, such as Chat GPT3.5, exhibited hallucination rates of 30-91% for academic citations^1,2^ and approximately 5-20% for code^3,4^, though more recent models have substantially improved on both^5^. However, the underlying probabilistic predictive process of LLMs, that gives rise to hallucinations, also enables LLMs to potentially predict beyond their training data; a capacity that in some respects may parallel human creativity in recombining known information in novel ways. The central question for any domain-specific application, such as metabolism studies, is therefore not whether hallucinations occur, but rather is whether the agent’s predictions can be computationally constrained so that this ability can be harnessed for predictive discovery. Further, all predictions must also be sufficiently interpretable and transparent such that expert users can distinguish plausible hypotheses from hallucination artifacts. Here we assess whether agentic AI has reached this threshold for drug metabolite annotation and discovery from mass spectrometry data. We are evaluating agentic AI not as a replacement for expert judgment, but as a tool that, when combined with domain expertise, can generate and prioritize candidate structures that would otherwise remain inaccessible.

Liquid chromatography tandem mass spectrometry (LC–MS/MS) is a central technology for the analysis of small molecules in metabolomics, natural products chemistry, and lipidomics, yet most detected ions remain structurally uncharacterized; typical spectral annotation rates are approximately 8% of detected features^6^. Structural interpretation requires integrating multiple sources of evidence. These include precursor *m/z*, isotopic patterns, MS/MS fragmentation, biochemical transformation rules, retention time, and biological context. The task of converging on a predicted structure is typically performed in a sequential, trial-and-error fashion by domain experts. Although computational tools exist for individual components of this process, including spectral similarity search (GNPS^7^, ModiFinder^8,9^, MS2Deepscore^10^, DreaMS^11^, ICEBERG^12,13^, molecular formula inference (SIRIUS^14^, BUDDY^15^, FIDDLE^16^), biotransformation analysis, and many others, they are generally applied independently and require substantial expertise to integrate into coherent structural hypotheses.

Repository-scale metabolomics infrastructures, including Metabolights^17^, Metabolomics Workbench^18^, GNPS/MassIVE^7^, have organized tandem mass spectrometry data, reference MS/MS spectra, and sample metadata within structured computational frameworks that enable systematic interrogation of chemical space across thousands of public studies using ReDU/panReDU^19,20^, MASST^21^, and MassQL^22^. The PanReDU alone indexes and harmonizes data from multiple metabolomics data repositories combining over 6,000 public metabolomics studies encompassing approximately 2 billion MS/MS spectra harmonized across 93 body sites, 25 health conditions, and 8,000 biological taxa. Despite this wealth of data, integrating these data resources and computational capabilities into coherent structural hypotheses remains a task that requires substantial expertise. We therefore assessed whether an agentic AI system could autonomously perform annotation and discovery from untargeted metabolomics data based on specific inquiries via data or via natural language.

## Results and discussion

### Agentic System Design

We integrated Claude Code (Anthropic; model claude-opus-4-6), an agentic AI system capable of autonomous multi-step reasoning, dynamic tool use, and iterative self-correction, as the backbone of a GNPS2 annotation and discovery pipeline. For each unknown/reference pair, the agent assembles spectral and chemical evidence to constrain its search space before reasoning begins, leveraging 26 domain-specific tools accessible through the Model Context Protocol (**SI Table 1 and methods**). Key tools include ModiFinder^8,9^ for MS/MS spectral alignment, which localizes the site of structural modification through shifted and unshifted fragment ion mapping; molecular formula decomposition of the observed mass difference; and a dual-layer candidate enumeration combining curated SMARTS-based transformation rules with SyGMa metabolic reaction rules^23^. Additional data resources integrated into the agent include a reference MS/MS library with SMILES annotations, panReDU sample level metadata, UniMod: a list of masses and explanations of post-translational modifications that can often be leveraged to also explain modifications in metabolomics data^24^, propagated MS/MS libraries including MassQL-generated libraries, MassQL query engine, MASST/FASST, extracted features, and FIDDLE: a molecular formula inference from MS/MS^16^. Ensuring all data sources shared a common format was essential to enabling AI ready agentic integration.

A critical design principle is that the agent is not constrained to select from pre-enumerated candidates. It can propose novel structures by reasoning over spectral evidence, writing and executing custom chemical analysis code, and drawing on biosynthetic and metabolic knowledge encoded in its tools and accompanying data. The agent maps observed fragment ions to proposed bond cleavages using MAGMa substructure annotations^25^, validates proposals through ICEBERG spectral prediction^13,26^, and assesses chemical plausibility (product stability, functional group compatibility, and metabolic reaction probability). Each prediction undergoes programmatic validation for mass accuracy, adduct consistency, and directional LogP-to-retention time agreement^27,28^ where retention time data are available. Validation failures trigger self-correction cycles in which the agent revises adduct assignments, re-evaluates modification sites, or generates alternative structures.

### Phase 1: Annotation Propagation of Drug Metabolite Candidates

To evaluate the agent’s annotation capability, and if it would provide any results that make sense and that we can confirm through creating synthetic standards, we analyzed 478 pairs of drugs and their putative structural analogs, i.e. a known drug and an unknown but putative structural analog guided by MS/MS similarity. These pairs were extracted from a drug incubation experiment examining microbial metabolism by a synthetic community of 111 bacterial species commonly found in the human gut^29,30^. These 478 pairs represent 73 unique known drugs and 424 unique but putative analogs, in-source fragments, multimers, or adducts (**SI Data 1**). To prioritize likely biotransformation products, we first agentically filtered pairs using the machine learning model STEP^31^ to prioritize single structural edits (filtered to a model score > 0.5) and selected the top 9 for agentic interpretation; for all 9 pairs, the agent produced structural predictions (**SI Data 2**). A second set of 30 pairs was prioritized by shifted peak count (≥5 shifted peaks, mass delta 16.1–300 Da, ≤3 pairs per drug); 19 of these 30 were annotated before a LLM token rate limit was reached on that session (**SI Data 5**). Out of the 28 explained examples, the agent reported that 14 are known and reported in PubChem but were not annotated by MS/MS library search. These include desmethoxyomeprazole derived from metabolism of omeprazole, both nitrile to amide conversion and further hydrolysis to the carboxylate of citalopram and demethylation of verapamil^32,33^.

We selected two of the previously unreported candidates for experimental validation in the biological samples based on synthetic accessibility and sample availability. As described in the LLM reports (**SI Data 3, 4, 6, and 7**), all the predictions and logic behind the structural proposals below were provided by the agentic AI without human involvement. From the first set of 9 pairs, an analog of hydroxyzine (antihistamine) was proposed by the agentic AI to have a mass delta of 79.9664 *m/z*, which it predicted to be matching to a phosphorylated species. (H_2_PO_4_, −0.9 ppm formula error; 0.03 ppm precursor mass error). The agent further identified the terminal primary hydroxyl of the 2-(2-hydroxyethoxy)ethyl chain as the sole plausible phosphorylation site, consistent with ModiFinder atom-level localization (**SI Data 2**); ether oxygens are non-nucleophilic and N-phosphorylation on tertiary amines is extremely rare. Thirteen unshifted fragment peaks confirmed the chlorophenyl-phenylmethyl-piperazine core was untouched, while the single shifted fragment (*m/z* 173→253, +79.97 Da) mapped directly to the hydroxyl-bearing terminus. All 22 agentic AI enumerated candidates converged on this structure (Agent Top-1 confidence: 0.92 - See methods for confidence guidance), and the agentic AI results described that the observed retention time decrease of −0.27 min was consistent with the addition of a polar phosphate group under reverse-phase conditions when compared to the parent drug. The agentic AI checked and according to the agentic AI, the predicted structure was not found in PubChem. Synthesis and LC-MS/MS confirmed the assignment as hydroxyzine phosphate, with MS/MS spectrum and retention time matching the biological sample when analyzed under identical conditions (**Fig. 1a– b**).

**Figure 1.**
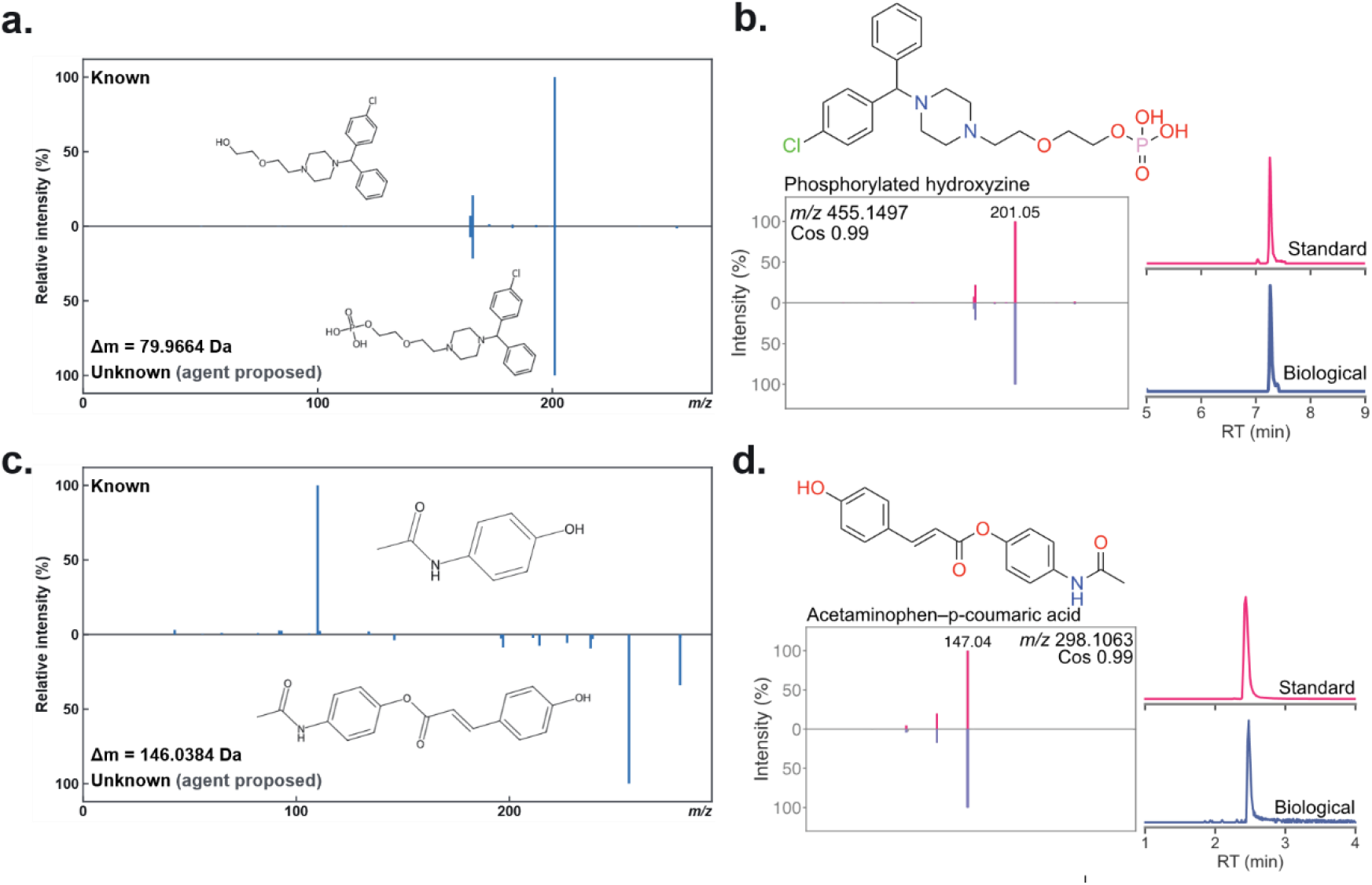
Agentic annotation propagation of drug metabolites. (a) MS/MS spectral alignment between hydroxyzine and its phosphorylated analog (Δ*m/z* 79.9664 Da; 0.03 ppm precursor mass error), with the agent-proposed phosphorylation at the terminal primary hydroxyl of the 2-(2-hydroxyethoxy)ethyl chain. The single shifted fragment (*m/z* 173→253) localizes the modification to the hydroxyl terminus; 13 unshifted peaks confirm the chlorophenyl-phenylmethyl-piperazine core is intact. (b) Confirmation by synthesis: MSlistic systems whose capacity to generate outputs beyond their training data, which can cause hallucinations, makes them also potentially suited to hypothesiz9H6O2, 5.4 ppm), with the agent-proposed acetaminophen–*p*-coumaric acid ester; all 5 shifted fragment peaks shift uniformly by ~146.04 Da. (d) Confirmation by synthesis: MS/MS spectrum and retention time match for acetaminophen–*p*-coumaric acid.

From the second set, an acetaminophen analog with a mass delta of 146.0384 *m/z* (C_9_H_6_O_2_, 5.4 ppm) was selected. The agent proposed ester bond formation between the phenol hydroxyl of acetaminophen and p-coumaric acid (loss of H_2_O), yielding C_17_H_15_NO_4_ (**See SI Data 6, 7 for logs**). All 5 shifted fragment peaks shifted uniformly by ~146.04 Da across substructures containing the full aromatic ring, consistent with the modification at the phenol oxygen. The large retention time increase of +3.71 min and LogP increase from 1.35 to 2.97 (+1.62) supported addition of a hydrophobic aromatic moiety rather than a sugar conjugate (which would decrease retention time). The agent ranked the p-coumarate ester (Agent Top-1 conf 0.55) above the m-coumarate (Agent confidence: 0.20) and o-coumarate isomers (Agent confidence: 0.15); p-coumaric acid is the most biosynthetically abundant hydroxycinnamic acid, and the o-isomer was further disfavored due to potential intramolecular cyclization to coumarin as described by the agent. Medium overall confidence was assigned because the ModiFinder match score was low and the specific positional isomer cannot be determined from mass alone. The predicted structure was not found in PubChem. It is important to note that this second set of 19 annotated pairs was incomplete; 11 additional pairs from the original 30-pair prioritization were not annotated due to our token limits being reached, and their structures remain uncharacterized. Synthesis and LC– MS/MS confirmed the p-coumarate assignment (**Figure 1c–d**). Although rare, phosphorylation as a conjugative biotransformation has been described for other drugs^34^, but to our knowledge no prior reports describe p-coumaric acid conjugation of a drug. The agent correctly identified a modification for which there was no prior example.

We note that not all agentic prediction tasks were successful, with some predictions being chemically implausible to be a metabolite but in many such cases the agent would provide caveats such as “The compound likely represents a synthetic analog rather than a metabolite.” or synthetically intractable. Nevertheless, the ability to semi-autonomously develop and test structural hypotheses substantially lowers the barrier to utilizing the many metabolomics tools that exist together with public data for discovery and just like decrease in unwanted hallucinations in coding and providing citations, we fully expect as agentic AI for structure predictions from mass spectrometry data will also improve.

### Phase 2: Repository-Scale Discovery of Undescribed Ibuprofen–Carnitine Conjugates

We recently established that ibuprofen and carnitine can be conjugated, resulting in delays in muscle recovery from birth injuries^35^. To determine whether public metabolomics repositories contained evidence of additional oxidative modifications of this conjugate, we prompted the GNPS2 agent with the natural-language query: “Are there carnitine conjugations of ibuprofen besides ibuprofen-carnitine, co-occurring in the same file as ibuprofen or ibuprofen-carnitine?” A full agentic AI report is available as supporting information (**SI Note 1, SI Data 8, 9**). The agent translated this query into a suspect screen across 3,604 datasets with MS/MS in GNPS2, we call Everything-Bagel corpus (**See methods**), using the diagnostic carnitine fragment at *m/z* 85.0284 and a mass-shifted ibuprofen acylium ion to confirm acylcarnitine class membership while localizing structural modifications to the ibuprofen moiety. Iterative refinement of mass accuracy tolerances and co-occurrence filtering against panReDU sample metadata identified seven candidate modification classes co-occurring with the ibuprofen–carnitine anchor, which was itself detected in 78 datasets.

The suspect screen identified 216 oxidative and carnitine conjugative candidate spectra across 26 datasets (40 of which the agent defined as high-confidence), all co-occurring in the same data files as the ibuprofen–carnitine anchor. Of particular note, 13 of the 78 datasets containing the ibuprofen–carnitine anchor were ibuprofen-named studies, meaning ibuprofen already appeared in the study title, sample metadata, or annotation table. All 78 datasets derive from human biofluids (urine, plasma, or feces), with the sole exception of a mouse body fluid study (MSV000101226), our own prior work demonstrating that ibuprofen slows muscle recovery from birth injuries and that this effect can be rescued by carnitine supplementation^35^, which motivated the present prompt. The two highest-confidence candidates that the agentic AI reported were hydroxy-ibuprofen-carnitine ([M+H]^+^ 366.2275, +O; detected in 11 datasets, 3 high-confidence) and carboxy-ibuprofen-carnitine ([M+H]^+^ 380.2068, +2O−2H; detected in 26 datasets, 24 high-confidence), the latter corresponding to the major urinary ibuprofen metabolite. Both were found in dataset accession MSV000099150. An oxidation ladder comprising dihydroxy (+2O, [M+H]^+^ 382.2224, 13 datasets), oxo (+O−2H, [M+H]^+^ 364.2118, 11 datasets), and trihydroxy (+3O, [M+H]^+^ 398.2173, 4 datasets) forms was additionally identified, with each member carrying the characteristic carnitine ion at *m/z* 85.0284 and an acylium ion mass-shifted by the added oxygens (**Figure 2, SI Data 9 and SI Table 2**). A hydroxymethyl (+CH_2_O, [M+H]^+^ 380.2431, 14 datasets) and glucuronide (+C_6_H_8_O_6_, [M+H]^+^ 526.2647, 13 datasets) form were also proposed but are treated as low-confidence leads: the agents reported that +CH_2_O delta is not a canonical ibuprofen metabolite, the glucuronide yielded zero high-confidence spectral matches, and mass deltas of ±CH_2_/±2H/+C_2_H_4_ are confounded by the endogenous acylcarnitine series. All of these assignments provided by the agent are at this stage hypothetical, being putative MS2-level annotations resting on anchor co-occurrence and the shifted acylium ion, with no authentic standards run, and none of the isomer positions unresolved (e.g., 2-OH vs. 3-OH), and specificity limited by the fact that the carnitine fragment at *m/z* 85.0284 is shared by all acylcarnitines. However, the majority of the predictions match known metabolism of ibuprofen itself^36^ but it was unknown they also potentially form carnitine conjugations. To test these hypotheses, hydroxy- and carboxy-ibuprofen-carnitine were selected for synthesis as the two highest-confidence candidates. In both cases the MS/MS spectra and retention times match the LC/MS data from the experimental samples (**Figure 2**). The remaining candidates and isomers await analogous synthetic confirmation.

**Figure 2.**
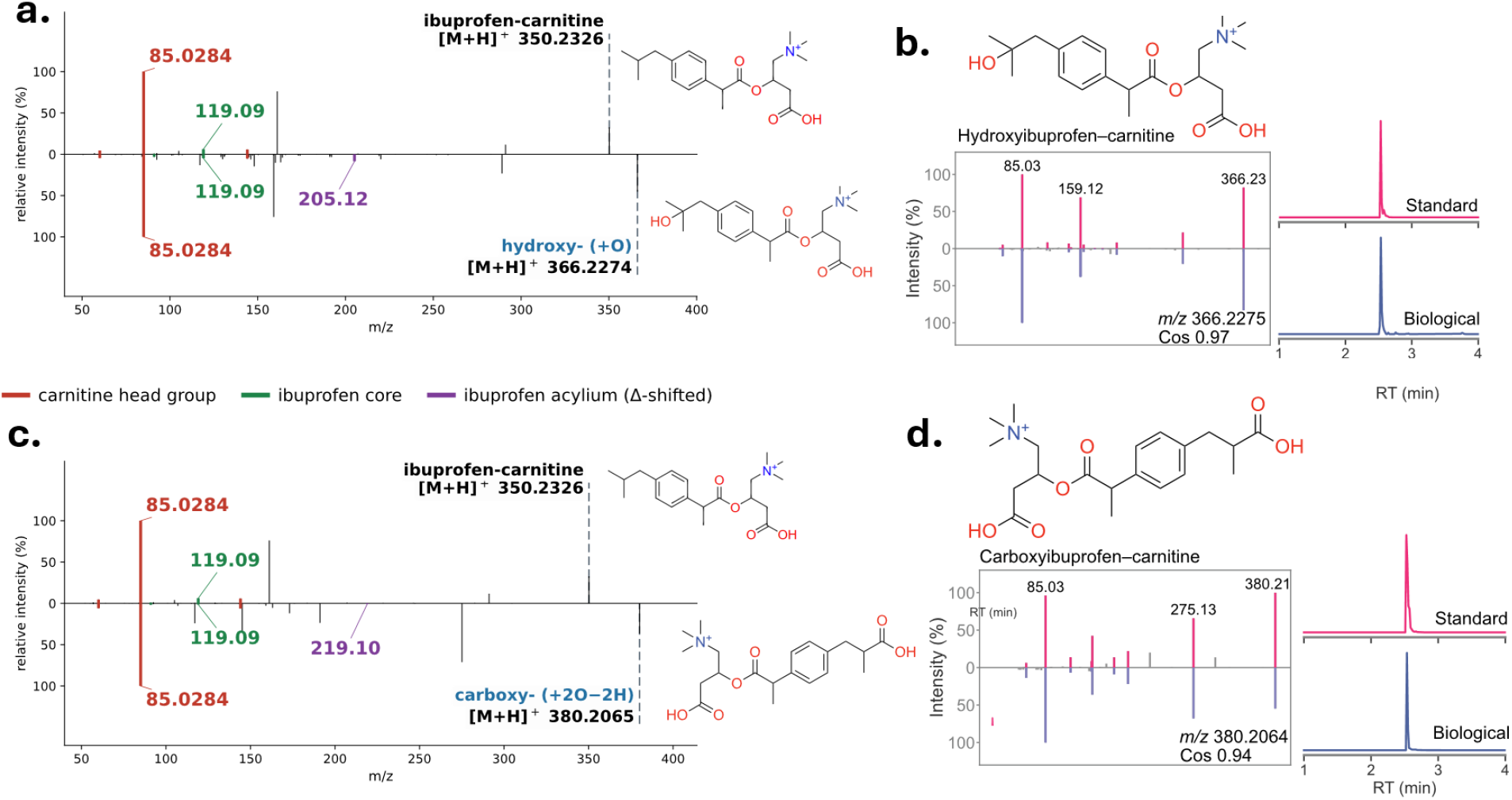
GNPS2 agent-guided discovery of ibuprofen–carnitine oxidative conjugates in public metabolomics datasets. (a) MS/MS spectral alignment of ibuprofen–carnitine ([M+H]^+^ 350.2326) and putative hydroxy-ibuprofen–carnitine ([M+H]^+^ 366.2275); the shifted acylium ion (+O, *m/z* 205.12) localizes the modification to the ibuprofen moiety while the carnitine ion (*m/z* 85.0284) confirms the acylcarnitine class. The GNPS2 agent acknowledged the ambiguity of 1, 2, or 3 hydroxyl isomers that could not be resolved. (b) Confirmation of the MS/MS and retention time match to hydroxy-ibuprofen–carnitine by matching against a synthetic standard. Note, we only synthesized the 1 position hydroxyl. As we did not synthesize all three isomers, we cannot exclude that 2 and 3 hydroxy isomers would also match the data. (c) MS/MS alignment for putative carboxy-ibuprofen–carnitine ([M+H]^+^ 380.2068); the shifted acylium (+2O-2H, *m/z* 219.10) localizes carboxylation to the ibuprofen moiety. (d) MS/MS and retention time match against a synthetic standard for carboxy-ibuprofen–carnitine.

## Conclusions

We present the GNPS2 agentic AI system as a proof-of-principle framework for autonomous structural elucidation and discovery in untargeted metabolomics. In two complementary applications, annotation propagation from known drug metabolites and repository-scale discovery of novel ibuprofen–carnitine conjugates, the system demonstrated that LLM-driven reasoning can integrate diverse computational tools and repository-scale MS/MS data to generate experimentally testable hypotheses for previously uncharacterized metabolites. We are explicit that the current system will produce incorrect predictions, and its full error rate cannot be exhaustively quantified at this stage; we therefore clearly frame this as a proof-of-principle demonstration whose accuracy will improve as tools, training data, reasoning strategies, and verification frameworks mature. We further note that repository-scale discovery queries across the Everything-Bagel corpus were unproductive if the queries were underspecified prompts and caused the agent to reach computational dead ends. This required multi-turn refinement of prompts to steer the agent toward productive tool use. In this proof of principle, we did not optimize for agentic efficiency or token usage in this first iteration nor address potential cybersecurity considerations. We currently addressed this by executing the agentic system in quarantined virtual machines which will require further development before broader deployment to the community.

## Acknowledgements

PCD acknowledges NIDDK R01DK136117, U24DK133658, and EnvedaGives Scientific Research Fund. MW acknowledges NIGMS 2R01GM107550. TRN and BPB acknowledge support from LBNL gift funds. HNZ acknowledges NIEHS 1K99ES037746.

## Methods

### Agentic AI for Structural Elucidation of Unknown Metabolites from Tandem Mass Spectrometry

We developed a set of skills and model context protocol (MCP) servers (**See SI Table 1 and Code Availability**) for the GNPS2 agent. All MCP servers were hosted in Docker that provided an interface to GNPS2 services and chemistry/metabolomics specific tools/data including FIDDLE, ICEBERG, ModiFinder, MASST, GNPS Libraries, UniMod, MiBiG, NPClassifier, SyGMa, and RDKit. We used Anthropic’s Claude Opus 4.6 in max effort large language model running natively in Claude Code on the Anthropic Max 20 subscription. This Claude Code was running on a Ubuntu Linux 20.04 virtual machine that had been isolated on a VLAN network with access to the internet. This virtual machine ran on a Dell Poweredge R730 with Intel Xeon CPU E5-2667 v3 @ 3.20GHz and 472.29 GB of RAM.

Our agentic annotation pipeline provided guidelines for ranking confidence of structural annotations to the LLM. First, for each analog structure candidate, we instructed the agent to provide a high/medium/low classification for the annotation on whether the known MS/MS and unknown MS/MS are structural analogs. For each predicted structure, the agent is guided to produce a top-1 structure confidence score that assesses compared to other isomers considered, how the agent scores the best candidate. This ranges between 0 and 1, with 1 being more confident.

### Everything-Bagel corpus construction

We collected public metabolomics data from public repositories, including MassIVE/GNPS, Metabolights, and Metabolomics Workbench into GNPS2. We chose only the LC/MS datasets that included MS/MS spectra. All LC/MS/MS were converted to open formats (mzXML and mzML) using msconvert [ref] with vendor centroiding of peaks. For each dataset, we performed a simple feature finding and alignment in order to de-duplicate MS/MS across each dataset. All MS/MS spectra were collected as MGF files for each dataset and collated together with one MGF per dataset with an additional attribution where each MS/MS and feature were found in the original files, with m/z and retention being key feature level metadata.

All sample level metadata across these dataset was downloaded from pan-ReDU (May 24, 2026) and joined by dataset and filename to each dataset MGF. This dataset formed the basis for the agent to conduct their discovery across 3,604 datasets.

### Drug Analogs Selection

Ten drug mixtures, with 14 drugs in each, were incubated with four defined microbial communities or a medium-only control and sampled at 0 and 72 h in triplicate. Samples were analyzed with LC-MS/MS in data-dependent acquisition mode, and feature extraction was performed with MZmine 4. To nominate candidate microbial drug metabolites, we filtered the feature table through successive abundance, temporal and annotation steps. We first retained features specific to a single drug mixture (in-versus out-mixture mean-abundance ratio > 10 at P < 0.01; one-sided Wilcoxon rank-sum test), then required these to be microbially dependent (culture-to-control mean ratio ≥ 3 at 72 h) and to accumulate over time (≥ 2-fold higher mean at 72 h than at 0 h). Surviving features were annotated as potential metabolites of incubated drugs based on MS/MS similarity calculated with both modified and reverse cosine (similarity score ≥ 0.7, matched peak ≥ 3). We finally retained potential drug metabolites whose retention time differs > 0.1 min from the parent drug to remove potential adducts or in-source fragments.

## Synthesis Methods

### Hydroxyzine phosphate

Hydroxyzine (0.50 mmol) was dissolved in anhydrous dichloromethane (DCM) and added dropwise to a stirred solution of phosphoryl chloride (POCl_3_, 0.60 mmol, 1.2 equiv) and triethylamine (TEA, 1.20 mmol, 2.4 equiv) in anhydrous DCM at (0 °C). After complete addition, the reaction mixture was allowed to warm gradually to room temperature and stirred for 24 h.

Deionized water (5 mL) was added slowly, and the resulting mixture was stirred for an additional 1 h. The solvent was removed under reduced pressure, and the residue was dissolved in DCM. The organic solution was washed with aqueous HCl (0.2 mM) and saturated brine three times. The organic layer was collected, dried over anhydrous MgSO_4_, and filtered. Removal of the solvent under reduced pressure afforded the crude product, which was washed with diethyl ether three times to yield the phosphorylated hydroxyzine derivative. The mixture was concentrated via rotary evaporator, after which the resulting crude product was subjected to purification on a CombiFlash NextGen 300+ system equipped with a 50 g Gold reverse-phase C18 column. Separation was carried out at a flow rate of 40 mL/min using water as Solvent A and ACN as Solvent B, following this gradient elution: 5% B held from 0 to 2 minutes; 5-10% B from 4 to 8 minutes; 40% B from 9 to 12 minutes; and 80% B from 13 to 16 minutes. Hydroxyzine phosphate fraction was collected during the 4–8 minute window at 5-10% B. Observed MS (+ESI): *m*/*z* = 455.1496 [M + H]^+^, C_21_H_28_ClN_2_O_5_P, (Exact calculated mass: 455.1497). ^1^H-NMR (MeOD) δ 3.56-3.64 (m, 8H), 3.71 (t, 2H), 3.89 (m, 6H), 5.64 (s, 1H), 7.42 (m, 1H), 7.49 (m, 4H), 7.88 (m, 4H). (The corresponding ^1^H-NMR spectrum is provided.) (^1^H-NMR spectra is available 10.5281/zenodo.20750582).

### Acetaminophen_*p*-coumaric acid

*p*-coumaric acid (1 equiv) and 2 mL of DMF were added to a 20 mL scintillation vial with a magnetic stir bar. To this solution, solid EDC (1.2 equiv) was subsequently added, and the solution was stirred at 23°C. After 15 minutes, acetaminophen 0.01 equiv, and DMAP (0.5 equiv) were added, and the reaction was stirred overnight. The mixture was concentrated via rotary evaporation, after which the resulting crude product was subjected to purification on a CombiFlash NextGen 300+ system equipped with a 25 g Gold normal-phase silica column. Separation was carried out at a flow rate of 25 mL/min using Dichloromethane as Solvent A and Methanol as Solvent B, following this gradient elution: 5% B held from 0 to 7 minutes; 20% B from 7 to 10 minutes; 40% B from 11 to 13 minutes; and 80% B from 13 to 15 minutes. The acetaminophen–p-coumaric acid ester fraction was collected during the 0–7 minute window at 5% B. Observed MS (+ESI): *m*/*z* = 298.1062 [M + H]^+^, C_17_H_15_NO_4_, (Exact calculated mass: 298.1074). ^1^H-NMR (MeOD) δ 2.15 (s, 3H), 6.53 (d, 1H), 6.85 (d, 2H), 7.13 (d, 2H), 7.55 (d, 2H), 7.80 (d, 1H). (The corresponding ^1^H-NMR spectrum is provided.) (^1^H-NMR spectra is available 10.5281/zenodo.20750582).

### Hydroxyibuprofen_carnitine and carboxyibuprofen_carnitine

Hydroxyibuprofen and carboxyibuprofen (1.0 equiv each) were dissolved separately in DMF (2 mL) in 20 mL scintillation vials equipped with magnetic stir bars. EDC.HCl (1.2 equiv) and DMAP (0.1 equiv) were added sequentially to each reaction mixture, and the resulting solutions were stirred at 23 °C for 15 min. Carnitine (1.2 equiv) was then added, and the reactions were stirred overnight at room temperature.

Upon completion, the reaction mixtures were concentrated under reduced pressure. The residues were dissolved in a suitable solvent and filtered prior to LC–MS/MS analysis. The resulting products were used as synthetic standards for retention-time matching with biological samples. Hydroxyibuprofen_carnitine: Observed MS (+ESI): *m*/*z* = 366.2275 [M]^+^, C_20_H_32_NO ^+^, (Exact calculated mass: 366.2275). Carboxyibuprofen_carnitine: Observed MS (+ESI): *m*/*z* = 380.2064 [M]^+^, C_20_H_30_NO ^+^, (Exact calculated mass: 380.2068).

Biological samples and synthetic standards were collected for retention time and MS/MS spectral matching and processed through LC-MS/MS analyses. These analyses were conducted on a Vanquish UHPLC system interfaced with a Q-Exactive Orbitrap mass spectrometer (Thermo Fisher Scientific, Bremen, Germany). Chromatographic separation was achieved on a Polar C18 column (Kinetex C18, 100 x 2.1 mm, 2.6 μm particle size, 100A pore size – Phenomenex, Torrance, USA), with a mobile phase composed of H_2_O (solvent A) and ACN (solvent B), each acidified with 0.1% formic acid. The following gradient was applied to assess retention time matching between the synthetic standards and the compounds detected in the biological samples: 0–0.5 min at 5% B; 0.5–1.1 min ramping from 5 to 30% B; 1.1–5.0 min ramping from 30 to 60% B; 5.0–9.0 min ramping from 60 to 100% B, followed by a 1.5 min wash at 100% B and a 1.5 min re-equilibration at 5% B. The flow rate was maintained at 0.5 mL/min, the injection volume was fixed at 3 μL, and the column temperature was held at 40 °C. Data-dependent acquisition (DDA) of MS/MS spectra was performed in positive ionization mode. Electrospray ionization (ESI) source parameters were configured as follows: sheath gas flow at 52.5 AU, auxiliary gas flow at 13.75 AU, spare gas flow at 2.7 AU, and auxiliary gas temperature at 400 °C; the spray voltage was adjusted to 3.5 kV, the inlet capillary temperature to 320 °C, and an S-lens level of 50 V was used. The MS scan range was defined as 300–800 m/z at a resolution of 35,000 with one micro-scan. The maximum ion injection time was set to 100 ms, with an automated gain control (AGC) target of 1.0E6. As many as 5 MS/MS spectra per MS1 survey scan were acquired in DDA mode at a resolution of 17,500 with one micro-scan. The maximum ion injection time for MS/MS scans was set to 150 ms, with an AGC target of 5E5 ions. The MS/MS precursor isolation window was defined as 1 m/z. Normalized collision energy was applied in a stepwise fashion at 30, 40, and 60, with a default charge state of z = 1. MS/MS scans were initiated at the apex of chromatographic peaks within a window of 2 to 5 seconds from their initial detection. Analytical quality and reproducibility were monitored by tracking the retention time and m/z values of a six-component standard mixture solution (amytriptiline, sulfamethizole, sulfamethoxine, sulfadimethoxine, coumarin 314, and sulfachlopyridazine), which was injected every 5 samples.

## Code Availability

Agentic Skills and MCP adapters can be found here: https://github.com/Wang-Bioinformatics-Lab/agent-annotation-plugin/blob/main/skill/SKILL.md

## Data Availability

Feature Based Molecular Networking of drug metabolism: https://gnps2.org/status?task=d24b86e7214c4888980a99059474befe

Dataset [MSV000099150](https://massive.ucsd.edu/ProteoSAFe/dataset.jsp?task=6e9d605d86f64f1cb3253ec8a1878e9f) is available publicly at MassIVE/GNPS.

Synthetic standards and biological samples Dataset [MSV000102198](https://massive.ucsd.edu/ProteoSAFe/dataset.jsp?task=a0a70a13eecd43b2af1e922143f87a75) is available publicly at MassIVE/GNPS.

