## Supplemental Information for "Agentic AI for Structural Elucidation and Discovery of Drug Metabolites from Mass Spectrometry Data"

**SUPPLEMENTARY INFORMATION**

**SI Data 1 - Candidate Drug Analogs Table**

**SI Data 2 - Top 9 molecules prioritized by STEP HTML visualization that prioritized - Prioritized Hydroxyzine**

**SI Data 3 - Hydroxyzine Phosphorylation Full Agentic LLM Log**

**SI Data 4 - Hydroxyzine Phosphorylation Readable Summary**

**SI Data 5 - Top 19 molecules prioritized by MS/MS matched peaks - Prioritized Acetaminophen**

**SI Data 6 - Acetaminophen and p-coumaric acid Full Agentic LLM Log**

**SI Data 7 - Acetaminophen and p-coumaric acid Readable Summary**

**SI Data 8 - Carnitine Repository Scale Search Full Agentic LLM Log**

**SI Data 9 - Carnitine Repository Scale LLM Summary Report**

**SI Table 1 - Domain-specific tools available to the GNPS2 Agent in Claude Code during execution**. Tools are accessible through the Model Context Protocol (MCP) and can be invoked autonomously by the GNPS2 agent for analog structure annotation.

| **Category** | **Tool** | **Parameters** | **Description** |
| --- | --- | --- | --- |
| Property Calculation | tool_calculate_logp | smiles | Calculates LogP (Wildman-Crippen), molecular weight, molecular formula, and hydrogen bond donor/acceptor counts |
|  | tool_identify_functional_groups | smiles | Identifies modifiable functional groups (tertiary amines, phenols, aromatic CH, hydroxyl, carboxyl, etc.) with atom indices |
|  | tool_describe_atom_environment | smiles, atom_indices | Describes the structural environment of specific atoms including element identity, ring membership, aromaticity, and nearby functional groups |
| Structure Validation | tool_validate_smiles | smiles, observed_mz, adduct | Validates SMILES syntax via RDKit and optionally checks mass error against observed precursor m/z |
|  | tool_detect_adduct | smiles, observed_mz | Tests 30+ adduct types against observed m/z to determine the most likely ionization form |
|  | tool_determine_molecular_formula | observed_mz, adduct, mass_tolerance_ppm | Determines possible molecular formulas from observed precursor m/z using MSBuddy mass decomposition |
|  | tool_compare_rt_logp | known_smiles, candidate_smiles, known_rt, unknown_rt | Checks whether the LogP change direction is consistent with the observed retention time shift (reverse-phase HPLC) |
| Structure Enumeration | tool_enumerate_modifications | known_smiles, delta_mass, modification_type, max_results | Enumerates candidate structures by applying SMARTS-based and SyGMa metabolic transformation rules |
| Structure Comparison | tool_compare_structures | smiles1, smiles2 | Computes Tanimoto similarity (Morgan fingerprints), maximum common substructure, and atom count difference |
| Spectral Analysis | tool_compare_spectra | peaks_a, peaks_b | Computes cosine similarity between two peak lists |
|  | tool_predict_spectrum | smiles, adduct | Predicts MS/MS spectrum using the ICEBERG neural network model |
|  | tool_predict_and_compare | candidate_smiles, experimental_peaks, adduct | Predicts MS/MS spectrum and computes cosine similarity against experimental spectrum |
|  | tool_fetch_spectrum_peaks | usi | Retrieves experimental MS/MS spectrum peaks by USI or local feature ID |
| Chemical Plausibility | tool_assess_plausibility | known_smiles, candidate_smiles, modification_type | Assesses plausibility based on product stability, functional group compatibility, and SyGMa metabolic probability |
| Compound Classification | tool_classify_compound | smiles | Classifies compounds by natural product pathway, superclass, and class using NPClassifier |
| Database Search | tool_search_known_compounds | smiles | Searches PubChem for exact structure match by SMILES |
|  | tool_check_known_compound | smiles | Combined PubChem and GNPS library search to determine if the candidate has been previously reported |
|  | tool_search_gnps_library | smiles, compound_name, precursor_mz, limit | Searches GNPS2 spectral library by SMILES, compound name, or precursor m/z |
|  | tool_search_fasst_library | usi, analog, min_cosine | Searches the GNPS FASST spectral library for similar reference spectra |
|  | tool_search_spectral_library | usi | Retrieves spectrum metadata from GNPS by USI |
| Modification Reference | tool_search_unimod | mass_difference, tolerance_da, top_k | Searches UniMod database for modifications matching an observed mass difference |
|  | tool_search_unimod_by_name | name, max_results | Searches UniMod modifications by name |
| Biosynthetic Knowledge | tool_search_mibig | compound, organism, biosynthesis_class | Searches MiBIG database for biosynthetic gene clusters |
|  | tool_get_biosynthetic_context | compound_class, known_smiles | Retrieves likely biotransformations for a compound class from the internal knowledge base |
| Formula Prediction | tool_predict_formula_from_spectrum | precursor_mz, peaks, adduct | Predicts molecular formula from MS/MS spectrum using the FIDDLE model |
| Code Execution | tool_run_chemistry_analysis | python_code, description | Executes arbitrary Python code with RDKit, NumPy, Pandas, SciPy |

**SI Note 1 Carnitine Conjugation Repository Scale Agent LLM Summary**

### **Phase 2 in Detail: Building and Running the Screen**

After the plan was approved ("ok lets give it a go"), Claude moved into implementation. This phase was the analytical heart of the project, and it's notable mostly for how many times the method had to be corrected as real data exposed flaws.

#### **The initial build and first smoke test**

Rather than writing the six separate scripts the plan envisioned, Claude consolidated the heavy lifting into a single parallel scan.py — reasoning that parsing ~34 GB of MGF spectral files was the bottleneck, so the corpus should be read only once. It then ran a smoke test on NORMAN (39 datasets) before committing to the full sweep.

That first run immediately exposed a problem: a ±0.02 Da mass window at m/z ~205 is about **96 ppm** — far too loose. Junk peaks were matching "ibuprofen," inflating the anchor counts.

#### **Three rounds of specificity tightening**

This is the most instructive part of Phase 2 — each fix came from looking at what the data actually returned rather than trusting the pipeline:

**Round 1 — tolerance and linkage logic.** Claude switched to a ppm-based precursor tolerance (15 ppm) and confronted a deeper issue: the carnitine diagnostic ion at m/z 85.0284 is shared by *every* acylcarnitine, so it alone can't tie a candidate to ibuprofen. The fix was to require two things: (a) co-occurrence with a confirmed ibuprofen-carnitine anchor in the same file, and (b) an **ibuprofen acylium fragment that is itself mass-shifted by the same modification delta** (e.g., a hydroxyl adds oxygen, shifting the acylium from 189.13 to 205.12).

**Round 2 — the homolog confound.** The next NORMAN run surfaced the subtlest issue. The top candidates were all +CH₂, +C₂H₄, and ±2H deltas — but those are *exactly the spacings of the natural fatty-acylcarnitine lipid series*. A "+CH₂ ibuprofen-carnitine" is mass-indistinguishable from simply the next ordinary acylcarnitine in the sample. Claude responded by classifying every delta into three buckets — **oxidative** (adding oxygen, which slides *off* the lipid series and is therefore specific), **conjugative** (phase-II additions, also specific), and **lipid_series** (the confounded ones) — and capped lipid-series hits at "low confidence" no matter how clean the evidence. It also required the anchor itself to look like ibuprofen (via aromatic-core fragments at 119/105/91), not just any C₂₀ acylcarnitine.

**Round 3 — fragment tolerance hygiene.** After inspecting real spectra, Claude noticed the m/z 144 carnitine ion was matching spuriously at −60 ppm because the 0.01 Da fragment window was too loose at higher masses. It tightened this to 0.006 Da and re-ran. The findings were essentially identical (78 anchors, ~399 candidates), which served as a robustness check.

#### **The full sweep and downstream stages**

With 32 cores available, Claude launched the full scan across 3,604 datasets using 24 workers (it ran in ~45 seconds in the final tuned version) and built the downstream stages while it ran:

- **Stage D** confirmed candidates co-occurred at the *same injection* level (shared non-zero peak area in the same raw run), not just the same dataset.
- **Stage E** joined results to ReDU sample metadata and study text, scoring biological plausibility — a drug metabolite is far more credible in human urine than in an environmental sample.

#### **Direct spectral validation**

This is where Claude earned the confidence in its result rather than just trusting mass matches. It manually inspected the actual peak lists:

- **MSV000099150** (human urine, study names ibuprofen): the anchor, hydroxy- (−0.3 ppm), and carboxy- (−0.7 ppm) conjugates all present, with the carnitine ion as base peak and the shifted acylium within a few ppm.
- **MSV000084744**: a full oxidation ladder — hydroxy → dihydroxy → trihydroxy → oxo — where the acylium fragment marched 205 → 221 → 237 (and 203 for oxo) with each added oxygen, while the aromatic core stayed put. This is exactly the fragmentation logic a real modified conjugate should show.
- **MSV000101226**: titled "Ibuprofen_carnitine_mouse_bodyfluid" — a purpose-built ibuprofen-carnitine study the pipeline recovered blind, functioning as a positive control.

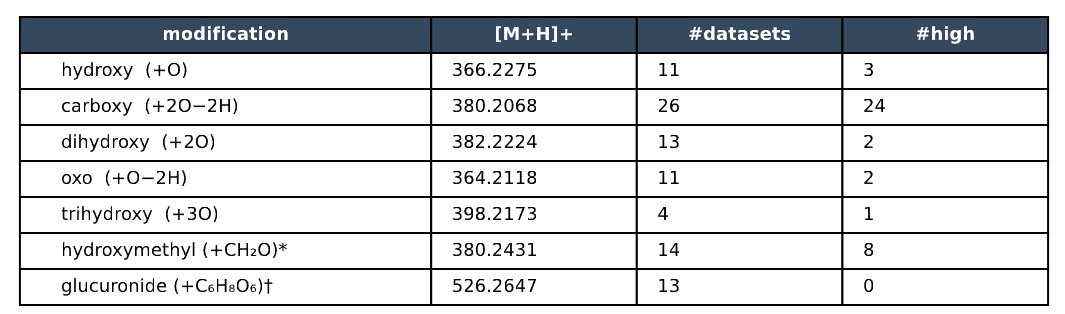

**SI Table 2** - Ibuprofen carnitines conjugates suggested by the GNPS2 agent in repository scale analysis, #high is number of MS/MS with proposed annotations the agent identified to be high scoring.
